# Candidate targets for immune responses to 2019-Novel Coronavirus (nCoV): sequence homology- and bioinformatic-based predictions

**DOI:** 10.1101/2020.02.12.946087

**Authors:** Alba Grifoni, John Sidney, Yun Zhang, Richard H Scheuermann, Bjoern Peters, Alessandro Sette

## Abstract

Effective countermeasures against the recent emergence and rapid expansion of the 2019-Novel Coronavirus (2019-nCoV) require the development of data and tools to understand and monitor viral spread and immune responses. However, little information about the targets of immune responses to 2019-nCoV is available. We used the Immune Epitope Database and Analysis Resource (IEDB) resource to catalog available data related to other coronaviruses, including SARS-CoV, which has high sequence similarity to 2019-nCoV, and is the best-characterized coronavirus in terms of epitope responses. We identified multiple specific regions in 2019-nCoV that have high homology to SARS virus. Parallel bionformatic predictions identified *a priori* potential B and T cell epitopes for 2019-nCoV. The independent identification of the same regions using two approaches reflects the high probability that these regions are targets for immune recognition of 2019-nCoV.

**ONE SENTENCE SUMMARY:** We identified potential targets for immune responses to 2019-nCoV and provide essential information for understanding human immune responses to this virus and evaluation of diagnostic and vaccine candidates.

## MAIN TEXT

On December 31, 2019, the Chinese Center for Disease Control (China CDC) reported a cluster of severe pneumonia cases of unknown etiology in the city of Wuhan in the Hubei province of China. Shortly thereafter, public health professionals identified the likely causative agent to be a novel *Betacoronavirus* (2019-nCoV). In collaboration with the China CDC and public health centers in other countries, the WHO reports 24,363 confirmed cases worldwide, with 491 deaths, as of February 5, 2020. Although the majority of cases have occurred in China, a small number have been confirmed in 24 other countries, including Japan, Thailand, South Korea, Singapore, Vietnam, India, the United States, Canada, Germany, France and the United Arab Emirates. These numbers are changing rapidly. For up-to-date information about the 2019-nCoV outbreak, see the WHO website at https://www.who.int/emergencies/diseases/novel-coronavirus-2019.

The Immune Epitope Database and Analysis Resource (IEDB) is a repository of epitope-related information curated from the scientific literature in the context of infectious disease, allergy and autoimmunity (*1*). The IEDB also provides bioinformatic tools and algorithms that allow for the analysis of epitope data and prediction of potential epitopes from novel sequences. The Virus Pathogen Resource (ViPR) is a complementary repository of information about human pathogenic viruses that integrates genome, gene, and protein sequence information with data about immune epitopes, protein structures, and host responses to virus infections (*2*).

Limited information is currently available about which parts of the 2019-nCoV sequence are recognized by human immune responses. Such knowledge is of immediate relevance, and would assist vaccine design and facilitate evaluation of vaccine candidate immunogenicity, as well as monitoring of the potential consequence of mutational events and epitope escape as the virus is transmitted through human populations.

Although no epitope data are yet available for 2019-nCoV, there is a significant body of information about epitopes for coronaviruses in general, and in particular for *Betacoronaviruses* like SARS-CoV and MERS-CoV which cause respiratory disease in humans (*3, 4*). Here, we used the IEDB and ViPR resources to compile known epitope sites from other coronaviruses, map corresponding regions in the 2019-nCoV sequences, and predict likely epitopes. We also used validated bioinformatic tools to predict B and T cell epitopes that are likely to be recognized in humans, and to assess the conservation of these epitopes across different coronavirus species.

Coronaviruses belong to the family *Coronaviradae*, order *Nidovirales*, and can be further subdivided into four main genera (*Alpha-, Beta-, Gamma*- and *Deltacoronaviruses*). Several *Alpha*- and *Betacoronaviruses* cause mild respiratory infections and common cold symptoms in humans, while others are zoonotic and infect birds, pigs, bats and other animals. In addition to 2019-nCoV, two other coronaviruses, SARS-CoV and MERS-CoV, caused large disease outbreaks that had high (10-30%) lethality rates and widespread societal impact upon emergence (**Fig 1**) (*3, 4*).

**Figure 1.**
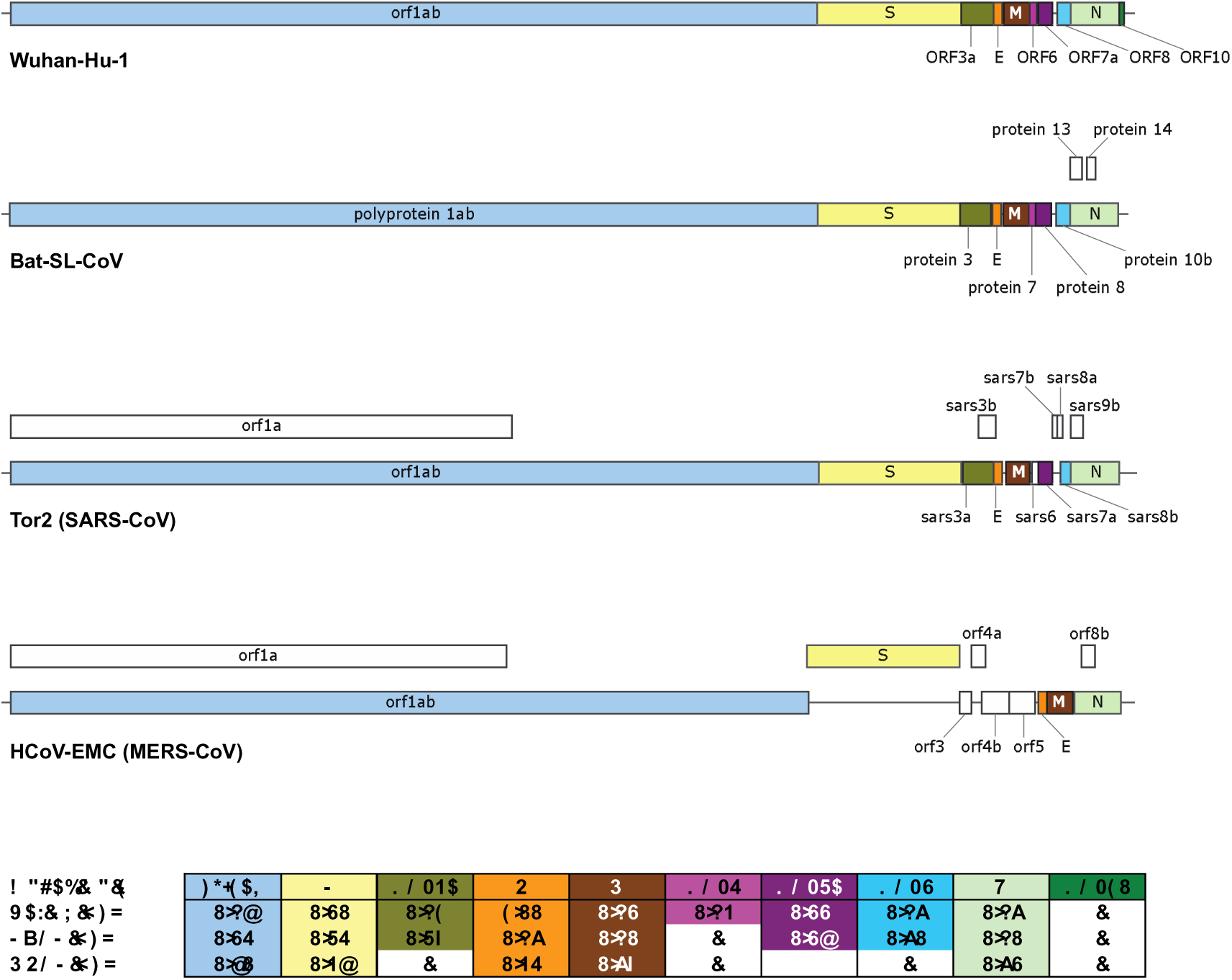
Comparison of 2019-nCoV (Wuhan-Hu-1) genome structure with its closest bat relative (bat-SL-CoVZXC21), Tor2 SARS-CoV and HCoV-EMC MERS-CoV. Above: CDS regions corresponding to homologous proteins between the four viruses are filled with the same color in the genome schematic to indicate homology; regions with no homology to the predicted 2019-nCoV proteins are colored white. Below: Table of pairwise protein similarities (expressed as % identity) between 2019-nCoV and the other three viruses.

The immune response to 2019-nCoV in humans awaits characterization, but human immune responses against other coronaviruses have been investigated. As of January 27, 2020, the IEDB has curated 581 linear, and 81 as discontinuous, B cell epitopes that have been reported in the peer reviewed literature. In addition, 320 peptides have been reported as T cell epitopes (**Table 1A**). The vast majority of these epitopes are derived from *Betacoronavirues*, and more specifically from SARS-CoV, which alone accounts for over 60% of them. In terms of the host in which the various B and T cell epitopes were recognized (**Table 1B)**, most epitopes (either B or T) were defined in humans or murine systems. Notably, all but two of the 417 B and T cell epitopes described in humans are from *Betacoronaviruses*, with 398 of them coming from SARS-CoV.

**Table 1.**
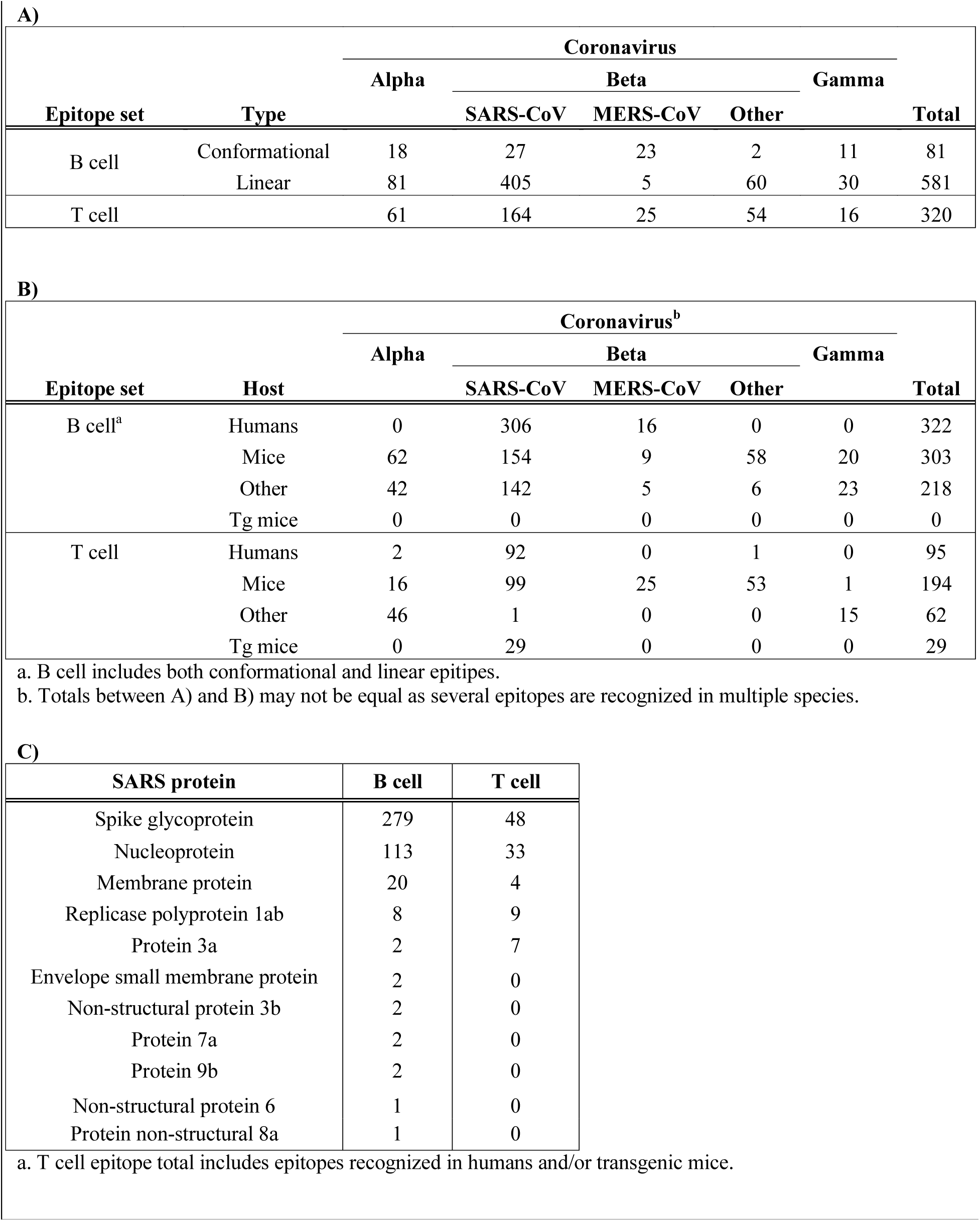
IEDB inventory of coronavirus B and T cell epitopes.

Comparison of a consensus 2019-nCoV protein sequence to sequences for SARS-CoV, MERS-CoV and bat-SL-CoVZXC21 revealed a high degree of similarity (expressed as % identity) between 2019-nCoV, bat-SL-CoVZXC21 and SARS-CoV, but more limited similarity with MERS-CoV (**Figure 1**). In conclusion, SARS-Cov is the closest related virus to 2019-nCoV for which a significant number of epitopes has been defined in humans (and other species), and that also causes human disease with lethal outcomes. Accordingly, in the following analyses we focused on comparing known SARS-CoV epitope sequences to the 2019-nCoV sequence.

We first assessed the distribution of SARS-derived epitopes as a function of protein of origin (**Table 1C)**. In the context of B cell responses, most of the 12 antigens in the SARS-CoV proteome are associated with epitopes, with the greatest number derived from spike glycoprotein, nucleoprotein and membrane protein (**Table 1C**). The paucity of B cell epitopes associated with the other proteins is likely because, on average, B cell epitope screening studies to date have probed regions constituting less than 20% of each respective sequence, including <1% of the Orf 1ab polyprotein. By comparison, the complete span of the spike glycoprotein, nucleoprotein and membrane protein sequences have been probed at least to some extent in B cell assays. A similar situation was observed in the case of T cell epitopes. Here we only considered epitopes whose recognition is restricted by human (HLA) MHC, since MHC polymorphism typically results in different epitopes being recognized in humans and mice.

B cell epitopes derived from SARS-CoV, were mapped back to a SARS-CoV reference sequence using the IEDB’s Immunobrowser tool (*5*). This tool combines all records available along a reference sequence and produces a Response Factor (RF) score that accounts for the positivity rate (how frequently a residue was found in a positive epitope) and the number of records (how many independent assays are reported). Dominant regions were identified considering residues stretches where the RF score was ≥0.3.

Analyses of the spike glycoprotein, membrane protein and nucleoproteins are shown in **Figure 2**. In the case of the spike glycoprotein (**Fig 2A)**, we identify five regions of potential interest (residues 274-306, 510-586, 587-628, 784-803 and 870-893), all representing regions associated with high immune response rates. Two regions were identified for membrane protein (1-25 and 131-152) (**Fig 2B)** and three regions were identified for nucleoprotein (43-65, 154-175 and 356-404) (**Fig 2C). Table 2** summarizes these analyses, identifying the specific regions associated with dominant B cell responses.

**Table 2.**
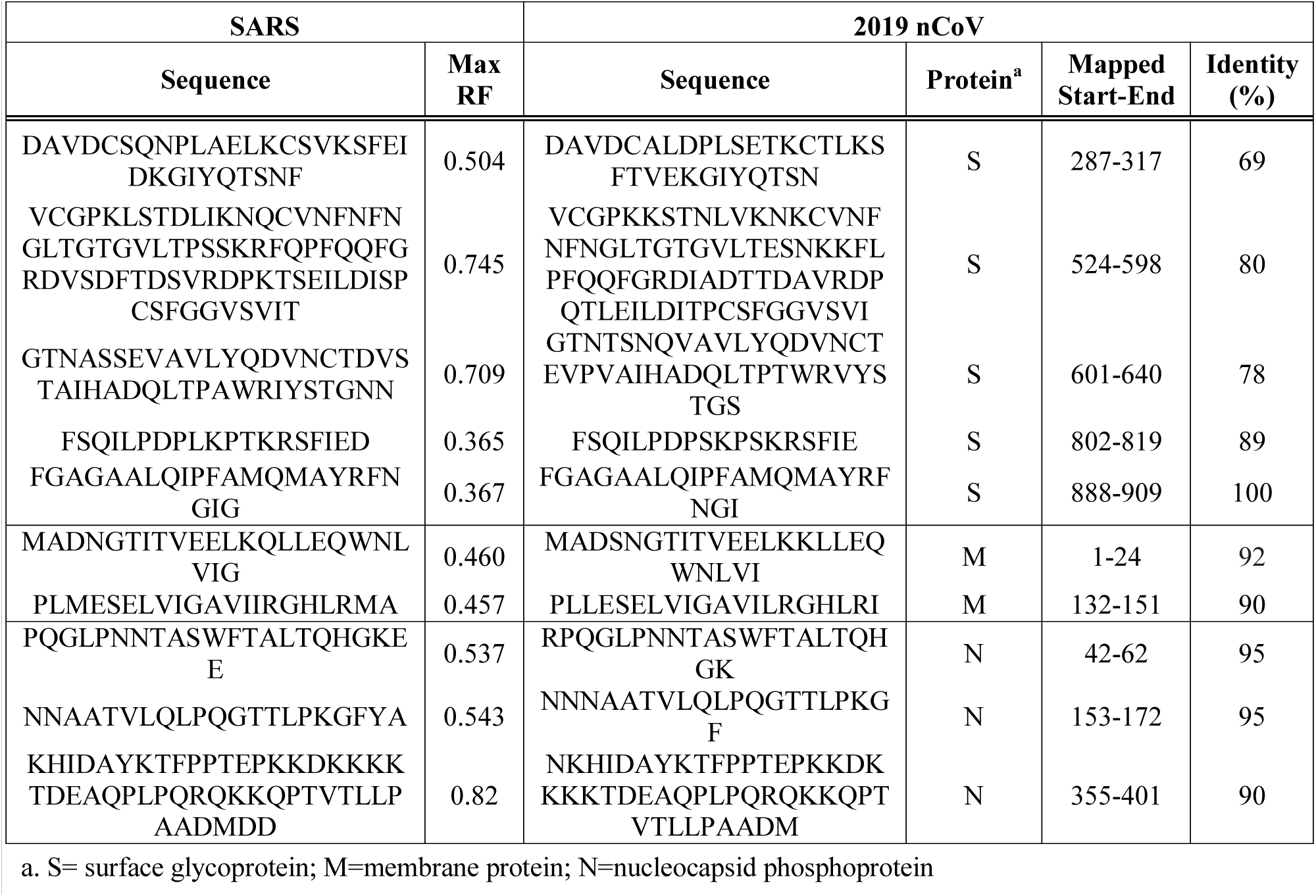
Dominant SARS-CoV B cell epitope regions.

**Figure 2.**
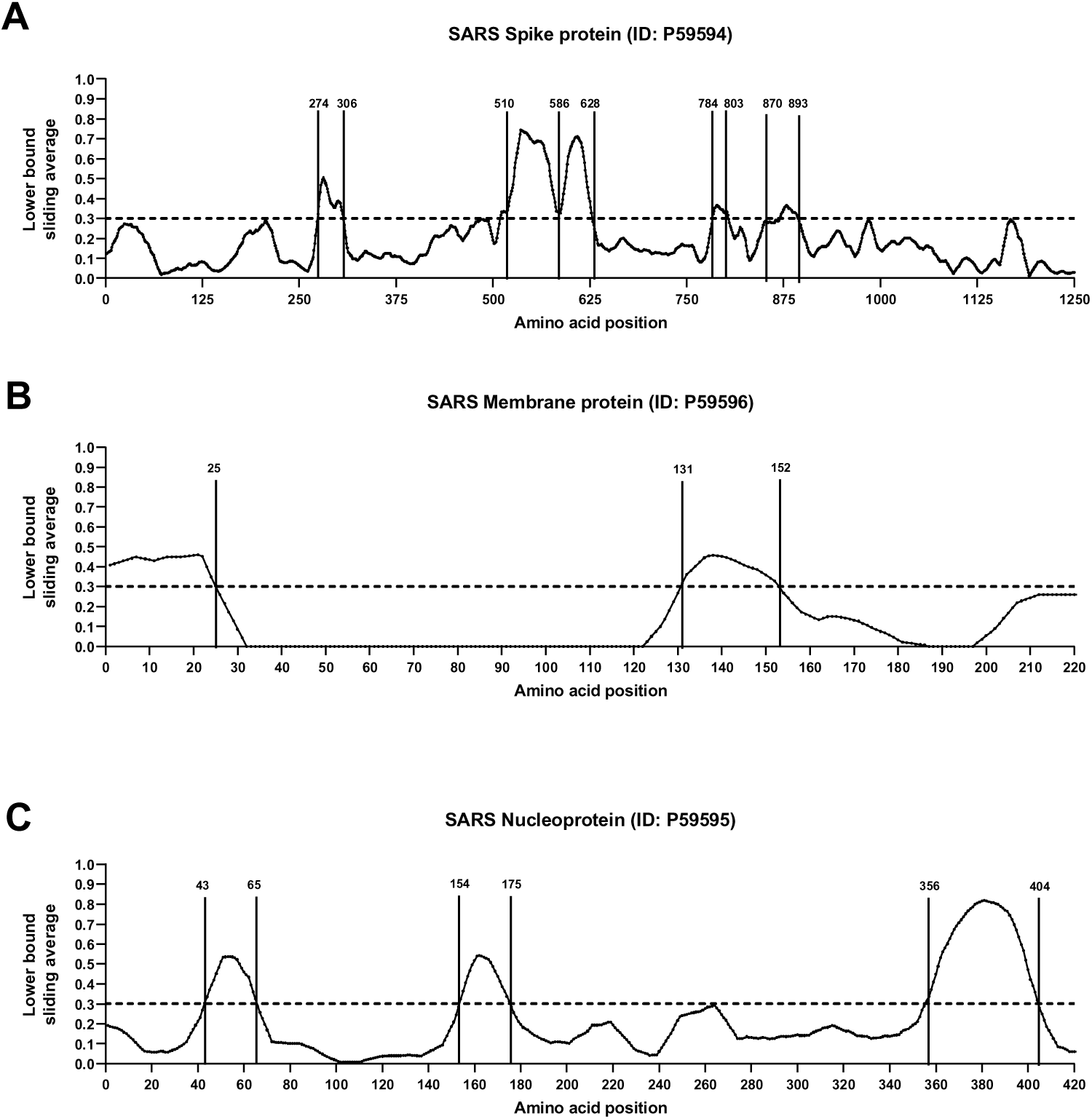
B cell immunodominant regions based on SARS-specific epitope mapping. RF score for each amino acid position was calculated (see Methods), and plotted over the SARS-CoV consensus sequence of spike glycoprotein (A), membrane protein (B) and nucleoprotein (C).

Next, we aligned the SARS-CoV B cell epitope region sequences to the 2019-nCoV sequence to calculate the percentage identity between each of the SARS-CoV dominant regions and 2019-nCoV (**Table 2**). Of the ten regions identified, six had 90% or more identity with 2019-nCoV, two were between 80-89% identical, and two had lower but still appreciable homology (69% and 78%). Because of the overall high level of sequence similarity of SARS-CoV and 2019-nCoV we infer that the same regions that are dominant in SARS-CoV have high likelihood to also be dominant in 2019-nCoV, even if the actual sequences are different.

In a similar analysis, T cell epitopes were also found to be predominantly associated with spike glycoprotein and nucleoprotein (**Table 1C**). However, in these cases, epitopic regions and individual epitopes were more widely dispersed throughout the respective proteins, which made identification of discrete, dominant epitopic regions more difficult. This outcome is not unexpected since T cells recognize short peptides generated from cellular processing of viral antigens that can be derived from any segment of the protein. **Table 3** shows a listing of the most dominant SARS-CoV individual epitopes identified to date in humans.

**Table 3.**
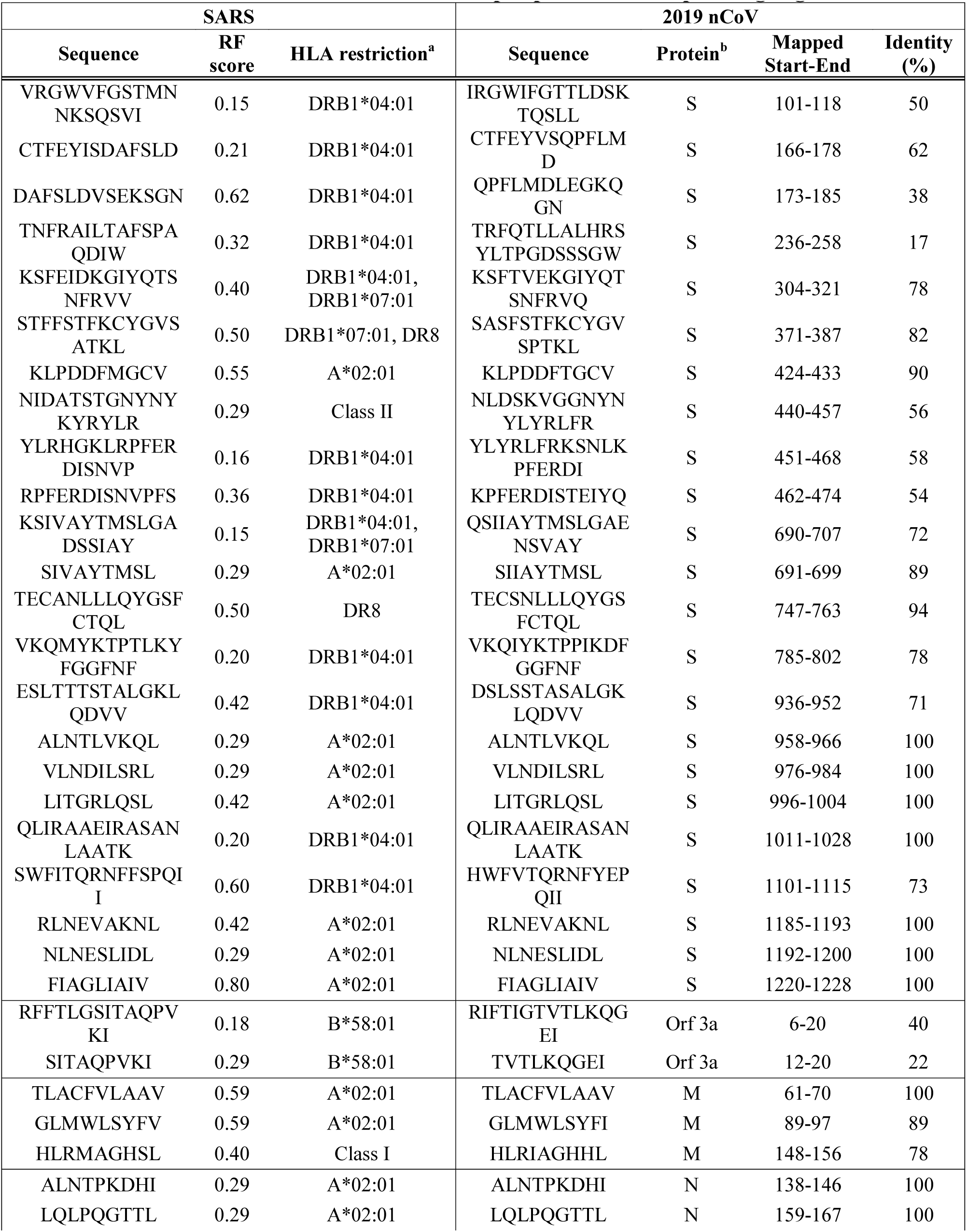
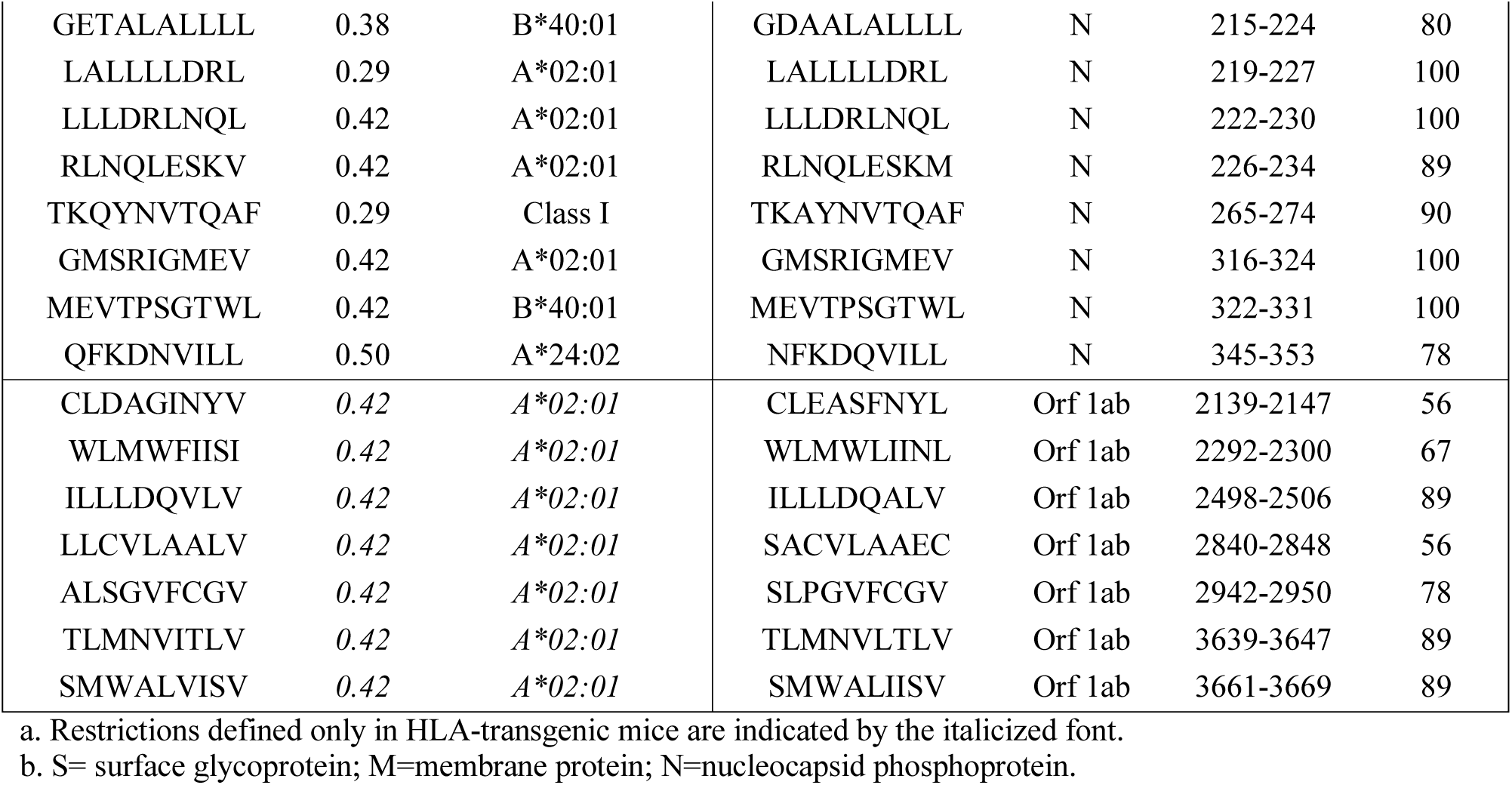
Dominant SARS-CoV-derived T cell epitopes and corresponding regions in nCoV.

We also aligned the SARS-CoV T cell epitope sequences and calculated for each epitope the percentage identity to 2019-nCoV. For each T cell epitope, **Table 3** shows the antigen of origin, the epitope sequence, the homologous 2019-nCoV sequence, and corresponding percentage of sequence identity. Overall, the nucleocapsid phosphoprotein and membrane-derived epitopes were most conserved (8/10 and 2/3, respectively, had ≥85% identity with 2019-nCoV). The Orf1ab and surface glycoprotein epitopes were moderately conserved (3/7 and 10/23, respectively, had ≥85% identity with 2019-nCoV), and Orf 3a epitopes were the least conserved.

To define potential B cell epitopes by an alternative method, we used the predictive tools provided with the IEDB Analysis Resource. B cell epitope predictions were carried out using the 2019-nCoV surface glycoprotein, nucleocapsid phosphoprotein, and membrane glycoprotein sequences, which, as described above, were found to be the main protein targets for B cell responses to other coronaviruses. In parallel, we performed predictions for linear B cell epitopes with Bepipred 2.0 (*6*), and for conformational epitopes with Discotope 2.0 (*7*). Both prediction algorithms are available on the IEDB B cell prediction tool page (http://tools.iedb.org/main/bcell/). A full list of B cell epitope prediction results per amino acid position per protein is provided in **Table S1**.

Using Bepipred 2.0 and a cutoff of ≥0.55 (corresponding to a specificity cutoff of 80%) (*6*), the surface glycoprotein had the highest number of predicted B cell epitopes, followed by membrane glycoprotein and nucleocapsid phosphoprotein (**Table S2**). To predict conformational B cell epitopes, we used SWISS-Model (*8*) and a SARS-CoV spike glycoprotein structure (PDB ID: 6ACD) to map the 2019-nCoV spike glycoprotein. No templates were found to build reliable models of the nucleocapsid phosphoprotein and membrane glycoprotein. A list of amino acid positions having a high probability of being included in predicted B cell epitopes for the surface glycoprotein, based on analysis with the Discotope 2.0 algorithm, are shown in **Table S1** (cutoff of ≥ -2.5 corresponding to 80% of specificity). We then localized the relevant amino acid positions in the 3D-model, and mapped them onto the model structure, which allowed identification of three relevant regions (444-449, 496-503, and 809-812) in the surface glycoprotein (**Figure 3**).

**Figure 3.**
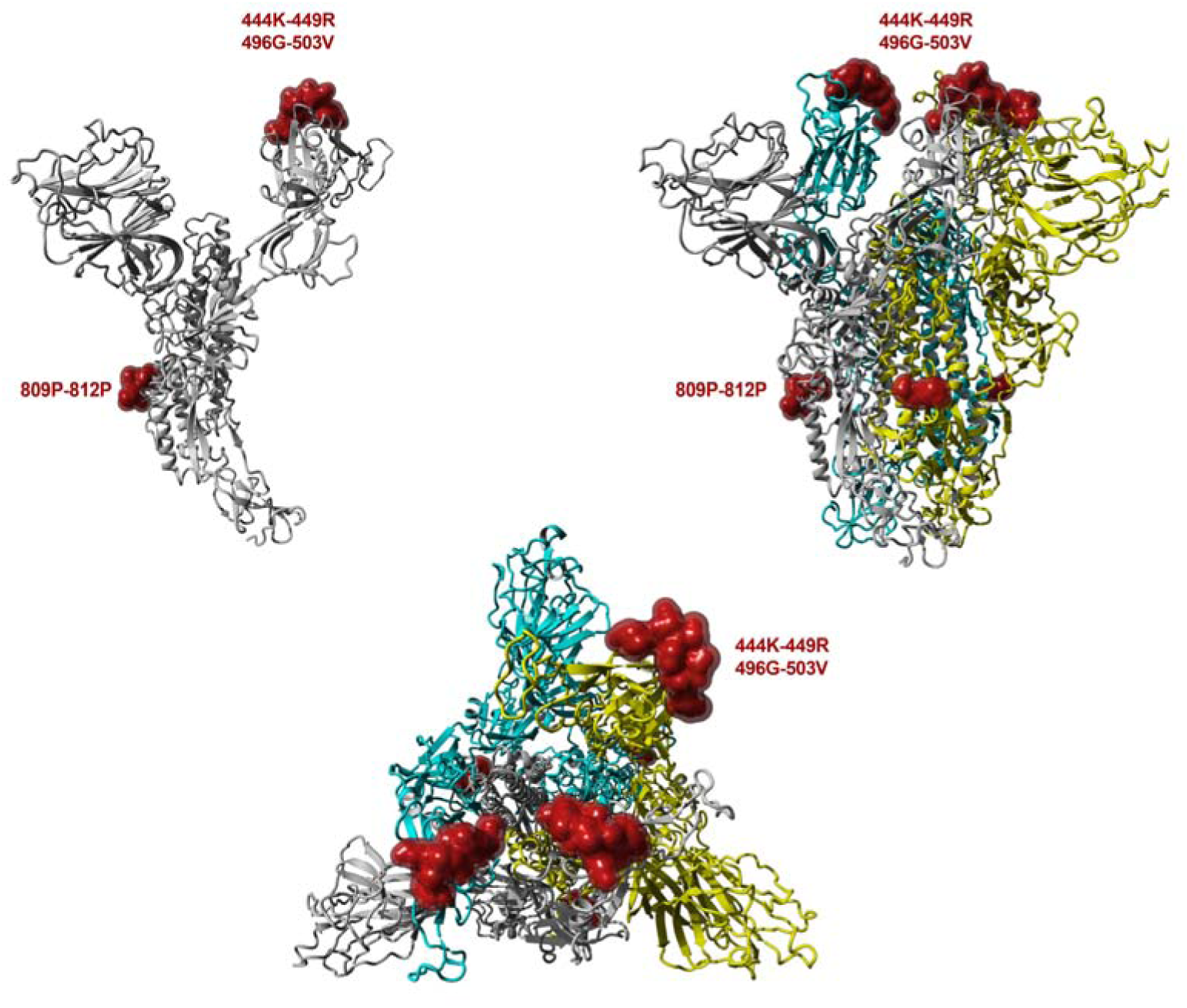
Model for 2019-nCoV surface glycoprotein structure based on the SARS-CoV spike glycoprotein structure (PDB: 6ACD). The calculated surface of the top 10 amino acid residues predicted to be B cell epitopes based on ranking performed with Discotope 2.0 are shown in red. The monomer is shown in the upper left, and the upper right and lower center present the trimer in two different orientations.

To predict CD4 T cell epitopes, we used the method described by Paul and co-authors (*9*), as implemented in the Tepitool resource in IEDB (*10*). This approach was designed and validated to predict dominant epitopes independently of ethnicity and HLA polymorphism, taking advantage of the extensive cross-reactivity and repertoire overlap between different HLA class II loci and allelic variants. Here, we selected peptides having a median consensus percentile ≤ 20, a threshold associated with epitope panels responsible for about 50% of target-specific responses. Using this threshold we identified 241 candidates in the 2019-nCoV sequence (see **Table S3**).

In previous experiments, we showed that pools based on similar peptide numbers can be generated by sequential lyophilization (*11*). These peptide pools (or megapools) incorporate predicted or experimentally-validated epitopes and allow measurement of magnitude and characterization of the phenotype of human T cell responses in infectious disease indications such as *Bordetella pertussis, Mycobacteria tuberculosis*, Dengue and Zika viruses (*11-14*). The 2019-nCoV CD4 megapool covers all 10 predicted proteins, with the number of potential epitopes proportional to the size of each protein (**Table S4**).

In parallel, we also sought to define likely CD8 epitopes. Here a different approach was required since the overlap between different HLA class I allelic variants and loci is more limited to specific groups of alleles, or supertypes (*15*). Following a previously validated approach (*16*), we assembled a set of the 12 most prominent HLA class I alleles which have been shown to allow broad coverage of the general population, as described in the Methods (see also **Table S5**). We then performed HLA class I binding predictions using the Net MHC pan 4.0 EL algorithm (*17*) available at the IEDB. For each allele, we selected the top 1% scoring peptides in the 2019-nCoV sequence, as ranked based on prediction. After eliminating redundancies and nested peptides, we obtained a final “*in silico*” megapool of 628 unique predicted epitopes. **Table S6** lists those unique predicted epitopes per protein, indicating for each their respective HLA restriction(s).

The epitopes identified by homology to the experimentally defined SARS-CoV epitopes shown in **Tables 2-3** were next compared with the epitopes identified by HLA-binding predicitons shown in **Tables S2-S3 and S6**. The epitopes independently identified in both approaches are presumed to be the most valuable leads. We first compared B cell immunodominant regions identified in SARS-CoV, and mapped to the homologous 2019-nCOV proteins (**Table 2**), with the predicted linear (**Table S2**) and conformational (**Table S1**) B cell epitopes. Out of the five B cell immunodominant regions from the SARS spike glycoprotein that were mapped to 2019-nCOV, three regions overlapped with those identified by BebiPred 2.0, and one overlapped with the 809-812 region predicted by Discotope 2.0 (**Table S1** and **Fig 3**). No overlap was observed for the five regions of SARS-CoV membrane protein and nucleoprotein that mapped to 2019-nCOV and those predicted by BebiPred 2.0. As stated above, no Discotope 2.0 prediction was available for those two proteins.

When we compared the SARS-CoV T cell epitopes that mapped to 2019-nCOV (**Table 3**) with the predicted CD4 and CD8 T cell epitopes (**Table S3** and **Table S6**, respectively), we found that 12 of 17 2019-nCOV T cell epitopes with high sequence identity (≥90%) to the SARS-CoV were independently identified by the two methods. Another 7 of 16 epitopes with moderate sequence identity (70-89%), and 6 of 12 epitopes with low sequence identity (<70%) were also identified by both methods. The lack of absolute correspondence is not surprising, given that the experimental data is derived from a skewed set of HLA restrictions (largely HLA A*02:01), and that our HLA class I prediction strategy targeted a more limited set of alleles that were selected to represent the most frequent worldwide variants; at the same time, the class II predictions are expected to cover 50% of the class II responses (*18*).

In conclusion, the use of available information related to SARS-CoV epitopes in conjunction with bionformatic predictions points to specific regions of 2019-nCoV which have a high likelihood of being recognized by human immune responses*. The observation that many B and T cell epitopes are highly conserved between 2019-nCoV and SARS-CoV is important. Protein regions that are conserved across relatively long evolutionary distances suggest that they are structurally or functionally constrained. Vaccination strategies designed to target the immune response toward these conserved epitope regions could generate immunity that is not only cross-protective across *Betacoronaviruses* but also relatively resistant to ongoing virus evolution.

* The corresponding peptide sets are being synthesized and will be made available for use by the scientific community upon request to the LJI team.

## METHODS

### IEDB analysis of T and B epitopes derived from coronaviruses

T and B cell epitopes for coronaviruses were identified by searching the IEDB at the end of January 2020. Queries were performed broadly for coronaviruses (taxonomy ID no. 11118), selecting positive assays in T cell, B cell and/or ligand contexts. Characteristics of each unique epitope (i.e., species, protein of provenance, positive assay type(s), MHC restriction) were tabulated, as well as the total number of donors tested and corresponding total number of donors with positive responses in B or T cell assays, and as a function of host. Finally, T or B cell assay specific response frequency scores (RF) were calculated broadly (i.e., any host), or for specific contexts (e.g., T cell assays in humans). Specifically, RF = [(r – sqrt(r)]/t, where r is the total number of responding donors and t is the total number of donors tested (*11*)).

SARS-CoV (tax ID no. 694009) sequence epitope density was visualized with the IEDB Immunobrowser tool (*5*). To identity contiguous dominant regions, RF scores for each residue were recalculated to represent a sliding 10 residue window.

### Selection of coronavirus proteome sequences for comparison to 2019-nCoV

All full-length protein sequences from SARS-CoV and MERS-CoV were retrieved from ViPR (https://www.viprbrc.org/brc/home.spg?decorator=corona) on 31 January 2020. In order to exclude sequences of experimental strains, sequences from “unknown,” mouse, and monkey hosts were excluded from analysis. Remaining sequences were aligned using the MUSCLE algorithm in ViPR. Sequences causing poor alignments in a preliminary analysis were removed before computing the final alignment. The consensus protein sequences of each virus group were determined from the final alignments using the Sequence Variation Analysis tool in ViPR. Protein sequences from natural virus isolates with sequences identical to the SARS-CoV and MERS-CoV consensus were selected for use in epitope sequence analysis.

### Determination of 2019-nCoV sequence conservation

Each Wuhan-Hu-1 (MN908947) protein sequence was compared against the consensus protein sequences from SARS-CoV and MERS-CoV and the protein sequences from closest bat relative (bat-SL-CoVZXC21) using the BLAST algorithm (ViPR; https://www.viprbrc.org/brc/blast.spg?method=ShowCleanInputPage&decorator=corona) to compute the pairwise identity between Wuhan-Hu-1 proteins and their comparison target.

### B cell epitope prediction on 2019-nCoV isolate

Linear B cell epitope predictions were carried out on three different coronavirus proteins: surface glycoprotein (S), nucleocapsid phosphoprotein (N) and membrane glycoprotein (M) (IDs: YP_009724390.1, YP_009724397.2 and YP_009724393.1, respectively) as the homologous versions of these proteins are the primary targets of B cell immune responses for SARS-CoV. We used the BebiPred 2.0(*6*) algorithm embedded in the B cell prediction analysis tool available in IEDB (*19*). For each protein, the epitope probability score for each amino acid and the probability of exposure was retrieved. Potential B cell epitopes were predicted using a cutoff of 0.55 (corresponding to specificity greater than 0.81 and sensitivity below 0.3) and considering sequences having more than 7 amino acid residues. Structure-based antibody prediction was performed by using Discotope 2.0 (*7*), available in IEDB (*19*) and a positivity cutoff greater than -2.5 was applied (corresponding to specificity greater than or equal to 0.80 and sensitivity below 0.39). Since no PDB structures are available in the Protein Data Bank (PDB) for the three 2019-nCoV proteins, a homology modeling approach was applied using SWISS-MODEL (*8*). Of the three proteins analyzed, only surface glycoprotein (S) had a PDB template that covered 0.93 of the entire protein based on the SARS-CoV spike glycoprotein (PDB ID: 6ACD) with a GMQE score of 0.72 and a QMEAN of -4.08; as such, it was the only model used for the structure-based B cell prediction. Additional information regarding the surface glycoprotein 2019-nCoV model are available in the SWISS-MODEL report (**Supplementary Fig S1)**.

### T cell epitope prediction on a 2019-nCoV isolate

Epitope prediction was carried out using the ten proteins predicted for the reference 2019-nCoV isolate, Wuhan-Hu-1. The corresponding protein accession identification numbers are: YP_009725255.1 (Orf 10), YP_009724397.2 (N), YP_009724396.1 (Orf 8), YP_009724395.1 (Orf 7a), YP_009724394.1 (Orf 6), YP_009724393.1 (M), YP_009724392.1 (Envelope protein, E), YP_009724391.1 (Orf 3a), YP_009724390.1 (S), and YP_009724389.1 (Orf 1ab).

For CD4 T cell epitope prediction, we applied a previously described algorithm that was developed to predict dominant HLA class II epitopes, using a median consensus percentile of prediction cutoff ≤20 as recommended (*18*). For CD8 T cell epitope prediction, we selected the 12 most frequent HLA class I alleles in the worldwide population (*20, 21*), using a phenotypic frequency cutoff ≥ 6%. The specific alleles included were: HLA-A*01:01, HLA-A*02:01, HLA-A*03:01, HLA-A*11:01, HLA-A*23:01, HLA-A*24:02, HLA-B*07:02, HLA-B*08:01, HLA-B*35:01, HLA-B*40:01, HLA-B*44:02, HLA-B*44:03. The 2019-nCoV protein sequences were run against this set of alleles using the NetMHCpan EL 4.0 algorithm and a size range of 8-14mers (*17*). For each HLA class I allele analyzed, we selected the top 1% epitopes ranked based on prediction score. To generate a final set for synthesis, duplicate peptides (i.e., those selected for multiple alleles) were reduced to a single occurance, and nested peptides were ensconced within longer sequences, up to 14 residues in length, before assigning the multiple corresponding HLA restrictions for each region.

## ACKNOWLEDGEMENTS

We thank Erica Ollmann Saphire, Sharon Schendel, and Mitchell Kronenberg for critical reading of the manuscript, and numerous helpful suggestions. Support for the work included funding provided through NIH-NIAID contracts 75N9301900065 (AS) and 75N93019C00001 (AS and BP). Additional support was provided through NIH-NIAID contract 75N93019C00076 (YZ and RS).

