## Supplementary material for "Candidate targets for immune responses to 2019-Novel Coronavirus (nCoV): sequence homology- and bioinformatic-based predictions": Figure S1

### SWISS-MODEL Homology Modelling Report

BIOZENTRUM University of Basel The Center for Molecular Life Sciences

#### Model Building Report

This document lists the results for the homology modelling project "2019 nCoV spike glycoprotein" submitted to SWISS-MODEL workspace on Jan. 29, 2020, 9:19 p.m..The submitted primary amino acid sequence is given in Table T1.

If you use any results in your research, please cite the relevant publications:


- Waterhouse, A., Bertoni, M., Bienert, S., Studer, G., Tauriello, G., Gumienny, R., Heer, F.T., de Beer, T.A.P., Rempfer, C., Bordoli, L., Lepore, R., Schwede, T. SWISS-MODEL: homology modelling of protein structures and complexes. Nucleic Acids Res. 46(W1), W296-W303 (2018). doi>
- Guex, N., Peitsch, M.C., Schwede, T. Automated comparative protein structure modeling with SWISS-MODEL and Swiss-PdbViewer: A historical perspective. Electrophoresis 30, S162-S173 (2009). doi>
- Bienert, S., Waterhouse, A., de Beer, T.A.P., Tauriello, G., Studer, G., Bordoli, L., Schwede, T. The SWISS-MODEL Repository - new features and functionality. Nucleic Acids Res. 45, D313-D319 (2017). doi>
- Benkert, P., Biasini, M., Schwede, T. Toward the estimation of the absolute quality of individual protein structure models. Bioinformatics 27, 343-350 (2011). doi>
- Bertoni, M., Kiefer, F., Biasini, M., Bordoli, L., Schwede, T. Modeling protein quaternary structure of homo- and hetero-oligomers beyond binary interactions by homology. Scientific Reports 7 (2017). doi>


#### Results

The SWISS-MODEL template library (SMTL version 2020-01-23, PDB release 2020-01-17) was searched with BLAST (Camacho et al.) and HHBlits (Remmert et al.) for evolutionary related structures matching the target sequence in Table T1. For details on the template search, see Materials and Methods. Overall 703 templates were found (Table T2).

#### Models

The following model was built (see Materials and Methods "Model Building"):

| Model #01 | File | Built with | Oligo-State | Ligands | GMQE | QMEAN |
|---|---|---|---|---|---|---|
| 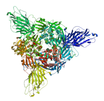 | PDB | ProMod3 2.0.0 | homo-trimer (matching prediction) | None | 0.72 | -4.08 |

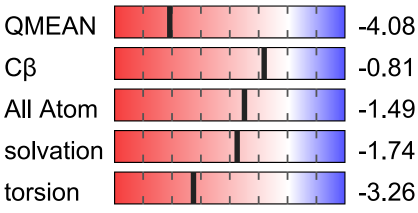


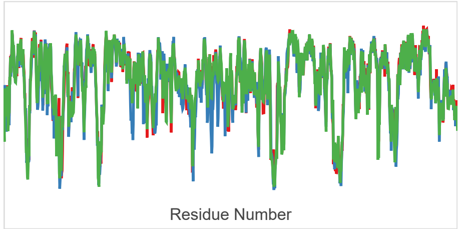


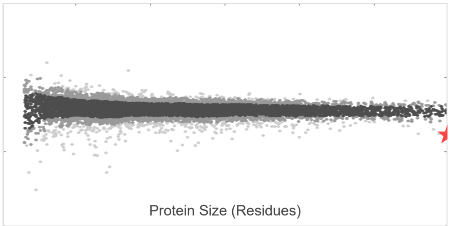


| Template | Seq Identity | Oligo-state | QSQE | Found by | Method | Resolution | Seq Similarity | Range | Coverage | Description |
|---|---|---|---|---|---|---|---|---|---|---|
| 6acd.1.A | 76.47 | homo-trimer | 0.75 | HHblits | EM | - | 0.54 | 15 - 1137 | 0.93 | Spike glycoprotein |

**The template contained no ligands.**

```
Target    MFVFLVLLP--LVSSQ--CVNLTTRTQLPPAYTNSFTRGVYYPDKVFRSSVLHSTQDLFLPFFSNVTWFHAIHVSGTNGT
6acd.1.A  MFIFLLFLTLTSGSDLDRCTTFDDVQAPNYTQHTSSMRGVYYPDEIFRSDTLYLTQDLFLPFYSNVTGFHTINH------

Target    KRFDNPVLPFNDGVYFASTEKSNIIRGWIFGTTLDSKTQSLLIVNNATNVVIKVCEFQFCNDPFLGVYY--HKNNKSWME
6acd.1.A  -TFGNPVIPFKDGIYFAATEKSNVVRGWVFGSTMNNKSQSVIIINNSTNVVIRACNFELCDNPFFAVSKPMGTQTHTM--
```

```
Target    SEFRVYSSANNCTFEYVSQPFLMDLEGKQGNFKNLREFVFKNIDGYFKIYSKHTPINLVRDLPQGFSALEPLVDLPIGIN
6acd.1.A  ----IFDNAFNCTFEYISDAFSLDVSEKSGNFKHLREFVFKNKDGFLYVYKGYQPIDVVRDLPSGFNTLKPIFKLPLGIN

Target    ITRFQTLLALHRSYLTPGDSSSGWTAGAAAYYVGYLQPRTFLLKYNENGTITDAVDCALDPLSETKCTLKSFTVEKGIYQ
6acd.1.A  ITNFRAILTAFSP----A--QDIWGTSAAAYFVGYLKPTTFMLKYDENGTITDAVDCSQNPLAELKCSVKSFEIDKGIYQ

Target    TSNFRVQPTESIVRFPNITNLCPFGEVFNATRFASVYAWNRKRISNCVADYSVLYNSASFSTFKCYGVSPTKLNDLCFTN
6acd.1.A  TSNFRVVPSGDVVRFPNITNLCPFGEVFNATKFPSVYAWERKKISNCVADYSVLYNSTFFSTFKCYGVSATKLNDLCFSN

Target    VYADSFVIRGDEVRQIAPGQTGKIADYNYKLPDDFTGCVIAWNSNNLDSKVGGNYNYLYRLFRKSNLKPFERDISTEIYQ
6acd.1.A  VYADSFVVKGDDVRQIAPGQTGVIADYNYKLPDDFMGCVLAWNTRNIDATSTGNYNYKYRYLRHGKLRPFERDISNVPFS

Target    AGSTPCNGVEGFNCYFPLQSYGFQPTNGVGYQPYRVVVLSFELLHAPATVCGPKKSTNLVKNKCVNFNFNGLTGTGVLTE
6acd.1.A  PDGKPCTP-PALNCYWPLNDYGFYTTTGIGYQPYRVVVLSFELLNAPATVCGPKLSTDLIKNQCVNFNFNGLTGTGVLTP

Target    SNKKFLPFQQFGRDIADTTDAVRDPQTLEILDITPCSFGGVSVITPGTNTSNQVAVLYQDVNCTEVPVAIHADQLTPTWR
6acd.1.A  SSKRFQPFQQFGRDVSDFTDSVRDPKTSEILDISPCSFGGVSVITPGTNASSEVAVLYQDVNCTDVSTAIHADQLTPAWR

Target    VYSTGSNVFQTRAGCLIGAEHVNNSYECDIPIGAGICASYQTQTNSPRRARSVASQSIIAYTMSLGAENSVAYSNNSIAI
6acd.1.A  IYSTGNNVFQTQAGCLIGAEHVDTSYECDIPIGAGICASYHTV----SLLRSTSQKSIVAYTMSLGADSSIAYSNNTIAI

Target    PTNFTISVTTEILPVSMTKTSVDCTMYICGDSTECSNLLLQYGSFCTQLNRALTGIAVEQDKNTQEVFAQVKQIYKTPPI
6acd.1.A  PTNFSISITTEVMPVSMAKTSVDCNMYICGDSTECANLLLQYGSFCTQLNRALSGIAAEQDRNTREVFAQVKQMYKTPTL

Target    KDFGGFNFSQILPDPSKPSKRSFIEDLLFNKVTLADAGFIKQYGDCLGDIAARDLICAQKFNGLTVLPPLLTDEMIAQYT
6acd.1.A  KYFGGFNFSQILPDPLKPTKRSFIEDLLFNKVTLADAGFMKQYGECLGDINARDLICAQKFNGLTVLPPLLTDDMIAAYT

Target    SALLAGTITSGWTFGAGAALQIPFAMQMAYRFNGIGVTQNVLYENQKLIANQFNSAIGKIQDSLSSTASALGKLQDVVNQ
6acd.1.A  AALVSGTATAGWTFGAGAALQIPFAMQMAYRFNGIGVTQNVLYENQKQIANQFNKAISQIQESLTTTSTALGKLQDVVNQ

Target    NAQALNTLVKQLSSNFGAISSVLNDILSRLDKVEAEVQIDRLITGRLQSLQTYVTQQLIRAAEIRASANLAATKMSECVL
6acd.1.A  NAQALNTLVKQLSSNFGAISSVLNDILSRLDKVEAEVQIDRLITGRLQSLQTYVTQQLIRAAEIRASANLAATKMSECVL

Target    GQSKRVDFCGKGYHLMSFPQSAPHGVVFLHVTYVPAQEKNFTTAPAICHDGKAHFPREGVFVSNGTHWFVTQRNFYEPQI
6acd.1.A  GQSKRVDFCGKGYHLMSFPQAAPHGVVFLHVTYVPSQERNFTTAPAICHEGKAYFPREGVFVFNGTSWFITQRNFFSPQI

Target    ITTDNTFVSGNCDVVIGIVNNTVYDPLQPELDSFKEELDKYFKNHTSPDVDLGDISGINASVVNIQKEIDRLNEVAKNLN
6acd.1.A  ITTDNTFVSGNCDVVIGIINNTVYDPLQPELDSFKEELDKYFKNHTSPDVDLGDISGINASVVNIQKEIDRLNEVAKNLN

Target    ESLIDLQELGKYEQYIKWPWYIWLGFIAGLIAIVMVTIMLCCMTSCCSCLKGCCSCGSCCKFDEDDSEPVLKGVKLHYT
6acd.1.A  ESLIDLQELGKYEQYIKWPW-----------------------------------------------------------


Target    MFVFLVLLP--LVSSQ--CVNLTTRTQLPPAYTNSFTRGVYYPDKVFRSSVLHSTQDLFLPFFSNVTWFHAIHVSGTNGT
6acd.1.C  MFIFLLFLTLTSGSDLDRCTTFDDVQAPNYTQHTSSMRGVYYPDEIFRSDTLYLTQDLFLPFYSNVTGFHTINH------

Target    KRFDNPVLPFNDGVYFASTEKSNIIRGWIFGTTLDSKTQSLLIVNNATNVVIKVCEFQFCNDPFLGVYY--HKNNKSWME
6acd.1.C  -TFGNPVIPFKDGIYFAATEKSNVVRGWVFGSTMNNKSQSVIIINNSTNVVIRACNFELCDNPFFAVSKPMGTQTHTM--

Target    SEFRVYSSANNCTFEYVSQPFLMDLEGKQGNFKNLREFVFKNIDGYFKIYSKHTPINLVRDLPQGFSALEPLVDLPIGIN
6acd.1.C  ----IFDNAFNCTFEYISDAFSLDVSEKSGNFKHLREFVFKNKDGFLYVYKGYQPIDVVRDLPSGFNTLKPIFKLPLGIN

Target    ITRFQTLLALHRSYLTPGDSSSGWTAGAAAYYVGYLQPRTFLLKYNENGTITDAVDCALDPLSETKCTLKSFTVEKGIYQ
6acd.1.C  ITNFRAILTAFSP----A--QDIWGTSAAAYFVGYLKPTTFMLKYDENGTITDAVDCSQNPLAELKCSVKSFEIDKGIYQ

Target    TSNFRVQPTESIVRFPNITNLCPFGEVFNATRFASVYAWNRKRISNCVADYSVLYNSASFSTFKCYGVSPTKLNDLCFTN
6acd.1.C  TSNFRVVPSGDVVRFPNITNLCPFGEVFNATKFPSVYAWERKKISNCVADYSVLYNSTFFSTFKCYGVSATKLNDLCFSN

Target    VYADSFVIRGDEVRQIAPGQTGKIADYNYKLPDDFTGCVIAWNSNNLDSKVGGNYNYLYRLFRKSNLKPFERDISTEIYQ
6acd.1.C  VYADSFVVKGDDVRQIAPGQTGVIADYNYKLPDDFMGCVLAWNTRNIDATSTGNYNYKYRYLRHGKLRPFERDISNVPFS

Target    AGSTPCNGVEGFNCYFPLQSYGFQPTNGVGYQPYRVVVLSFELLHAPATVCGPKKSTNLVKNKCVNFNFNGLTGTGVLTE
6acd.1.C  PDGKPCTP-PALNCYWPLNDYGFYTTTGIGYQPYRVVVLSFELLNAPATVCGPKLSTDLIKNQCVNFNFNGLTGTGVLTP
```

```
Target    SNKKFLPFQQFGRDIADTTDAVRDPQTLEILDITPCSFGGVSVITPGTNTSNQVAVLYQDVNCTEVPVAIHADQLTPTWR
6acd.1.C  SSKRFQPFQQFGRDVSDFTDSVRDPKTSEILDISPCSFGGVSVITPGTNASSEVAVLYQDVNCTDVSTAIHADQLTPAWR

Target    VYSTGSNVFQTRAGCLIGAEHVNNSYECDIPIGAGICASYQTQTNSPRRARSVASQSIIAYTMSLGAENSVAYSNNSIAI
6acd.1.C  IYSTGNNVFQTQAGCLIGAEHVDTSYECDIPIGAGICASYHTV----SLLRSTSQKSIVAYTMSLGADSSIAYSNNTIAI

Target    PTNFTISVTTEILPVSMTKTSVDCTMYICGDSTECSNLLLQYGSFCTQLNRALTGIAVEQDKNTQEVFAQVKQIYKTPPI
6acd.1.C  PTNFSISITTEVMPVSMAKTSVDCNMYICGDSTECANLLLQYGSFCTQLNRALSGIAAEQDRNTREVFAQVKQMYKTPTL

Target    KDFGGFNFSQILPDPSKPSKRSFIEDLLFNKVTLADAGFIKQYGDCLGDIAARDLICAQKFNGLTVLPPLLTDEMIAQYT
6acd.1.C  KYFGGFNFSQILPDPLKPTKRSFIEDLLFNKVTLADAGFMKQYGECLGDINARDLICAQKFNGLTVLPPLLTDDMIAAYT

Target    SALLAGTITSGWTFGAGAALQIPFAMQMAYRFNGIGVTQNVLYENQKLIANQFNSAIGKIQDSLSSTASALGKLQDVVNQ
6acd.1.C  AALVSGTATAGWTFGAGAALQIPFAMQMAYRFNGIGVTQNVLYENQKQIANQFNKAISQIQESLTTTSTALGKLQDVVNQ

Target    NAQALNTLVKQLSSNFGAISSVLNDILSRLDKVEAEVQIDRLITGRLQSLQTYVTQQLIRAAEIRASANLAATKMSECVL
6acd.1.C  NAQALNTLVKQLSSNFGAISSVLNDILSRLDKVEAEVQIDRLITGRLQSLQTYVTQQLIRAAEIRASANLAATKMSECVL

Target    GQSKRVDFCGKGYHLMSFPQSAPHGVVFLHVTYVPAQEKNFTTAPAICHDGKAHFPREGVFVSNGTHWFVTQRNFYEPQI
6acd.1.C  GQSKRVDFCGKGYHLMSFPQAAPHGVVFLHVTYVPSQERNFTTAPAICHEGKAYFPREGVFVFNGTSWFITQRNFFSPQI

Target    ITTDNTFVSGNCDVVIGIVNNTVYDPLQPELDSFKEELDKYFKNHTSPDVDLGDISGINASVVNIQKEIDRLNEVAKNLN
6acd.1.C  ITTDNTFVSGNCDVVIGIINNTVYDPLQPELDSFKEELDKYFKNHTSPDVDLGDISGINASVVNIQKEIDRLNEVAKNLN

Target    ESLIDLQELGKYEQYIKWPWYIWLGFIAGLIAIVMVTIMLCCMTSCCSCLKGCCSCGSCCKFDEDDSEPVLKGVKLHYT
6acd.1.C  ESLIDLQELGKYEQYIKWPW-----------------------------------------------------------


Target    MFVFLVLLP--LVSSQ--CVNLTTRTQLPPAYTNSFTRGVYYPDKVFRSSVLHSTQDLFLPFFSNVTWFHAIHVSGTNGT
6acd.1.B  MFIFLLFLTLTSGSDLDRCTTFDDVQAPNYTQHTSSMRGVYYPDEIFRSDTLYLTQDLFLPFYSNVTGFHTINH------

Target    KRFDNPVLPFNDGVYFASTEKSNIIRGWIFGTTLDSKTQSLLIVNNATNVVIKVCEFQFCNDPFLGVYY--HKNNKSWME
6acd.1.B  -TFGNPVIPFKDGIYFAATEKSNVVRGWVFGSTMNNKSQSVIIINNSTNVVIRACNFELCDNPFFAVSKPMGTQTHTM--

Target    SEFRVYSSANNCTFEYVSQPFLMDLEGKQGNFKNLREFVFKNIDGYFKIYSKHTPINLVRDLPQGFSALEPLVDLPIGIN
6acd.1.B  ----IFDNAFNCTFEYISDAFSLDVSEKSGNFKHLREFVFKNKDGFLYVYKGYQPIDVVRDLPSGFNTLKPIFKLPLGIN

Target    ITRFQTLLALHRSYLTPGDSSSGWTAGAAAYYVGYLQPRTFLLKYNENGTITDAVDCALDPLSETKCTLKSFTVEKGIYQ
6acd.1.B  ITNFRAILTAFSP----A--QDIWGTSAAAYFVGYLKPTTFMLKYDENGTITDAVDCSQNPLAELKCSVKSFEIDKGIYQ

Target    TSNFRVQPTESIVRFPNITNLCPFGEVFNATRFASVYAWNRKRISNCVADYSVLYNSASFSTFKCYGVSPTKLNDLCFTN
6acd.1.B  TSNFRVVPSGDVVRFPNITNLCPFGEVFNATKFPSVYAWERKKISNCVADYSVLYNSTFFSTFKCYGVSATKLNDLCFSN

Target    VYADSFVIRGDEVRQIAPGQTGKIADYNYKLPDDFTGCVIAWNSNNLDSKVGGNYNYLYRLFRKSNLKPFERDISTEIYQ
6acd.1.B  VYADSFVVKGDDVRQIAPGQTGVIADYNYKLPDDFMGCVLAWNTRNIDATSTGNYNYKYRYLRHGKLRPFERDISNVPFS

Target    AGSTPCNGVEGFNCYFPLQSYGFQPTNGVGYQPYRVVVLSFELLHAPATVCGPKKSTNLVKNKCVNFNFNGLTGTGVLTE
6acd.1.B  PDGKPCTP-PALNCYWPLNDYGFYTTTGIGYQPYRVVVLSFELLNAPATVCGPKLSTDLIKNQCVNFNFNGLTGTGVLTP

Target    SNKKFLPFQQFGRDIADTTDAVRDPQTLEILDITPCSFGGVSVITPGTNTSNQVAVLYQDVNCTEVPVAIHADQLTPTWR
6acd.1.B  SSKRFQPFQQFGRDVSDFTDSVRDPKTSEILDISPCSFGGVSVITPGTNASSEVAVLYQDVNCTDVSTAIHADQLTPAWR

Target    VYSTGSNVFQTRAGCLIGAEHVNNSYECDIPIGAGICASYQTQTNSPRRARSVASQSIIAYTMSLGAENSVAYSNNSIAI
6acd.1.B  IYSTGNNVFQTQAGCLIGAEHVDTSYECDIPIGAGICASYHTV----SLLRSTSQKSIVAYTMSLGADSSIAYSNNTIAI

Target    PTNFTISVTTEILPVSMTKTSVDCTMYICGDSTECSNLLLQYGSFCTQLNRALTGIAVEQDKNTQEVFAQVKQIYKTPPI
6acd.1.B  PTNFSISITTEVMPVSMAKTSVDCNMYICGDSTECANLLLQYGSFCTQLNRALSGIAAEQDRNTREVFAQVKQMYKTPTL

Target    KDFGGFNFSQILPDPSKPSKRSFIEDLLFNKVTLADAGFIKQYGDCLGDIAARDLICAQKFNGLTVLPPLLTDEMIAQYT
6acd.1.B  KYFGGFNFSQILPDPLKPTKRSFIEDLLFNKVTLADAGFMKQYGECLGDINARDLICAQKFNGLTVLPPLLTDDMIAAYT

Target    SALLAGTITSGWTFGAGAALQIPFAMQMAYRFNGIGVTQNVLYENQKLIANQFNSAIGKIQDSLSSTASALGKLQDVVNQ
```

```
6acd.1.B  AALVSGTATAGWTFGAGAALQIPFAMQMAYRFNGIGVTQNVLYENQKQIANQFNKAISQIQESLTTTSTALGKLQDVVNQ

Target    NAQALNTLVKQLSSNFGAISSVLNDILSRLDKVEAEVQIDRLITGRLQSLQTYVTQQLIRAAEIRASANLAATKMSECVL
6acd.1.B  NAQALNTLVKQLSSNFGAISSVLNDILSRLDKVEAEVQIDRLITGRLQSLQTYVTQQLIRAAEIRASANLAATKMSECVL

Target    GQSKRVDFCGKGYHLMSFPQSAPHGVVFLHVTYVPAQEKNFTTAPAICHDGKAHFPREGVFVSNGTHWFVTQRNFYEPQI
6acd.1.B  GQSKRVDFCGKGYHLMSFPQAAPHGVVFLHVTYVPSQERNFTTAPAICHEGKAYFPREGVFVFNGTSWFITQRNFFSPQI

Target    ITTDNTFVSGNCDVVIGIVNNTVYDPLQPELDSFKEELDKYFKNHTSPDVDLGDISGINASVVNIQKEIDRLNEVAKNLN
6acd.1.B  ITTDNTFVSGNCDVVIGIINNTVYDPLQPELDSFKEELDKYFKNHTSPDVDLGDISGINASVVNIQKEIDRLNEVAKNLN

Target    ESLIDLQELGKYEQYIKWPWYIWLGFIAGLIAIVMVTIMLCCMTSCCSCLKGCCSCGSCCKFDEDDSEPVLKGVKLHYT
6acd.1.B  ESLIDLQELGKYEQYIKWPW-----------------------------------------------------------
```

---

#### Materials and Methods

##### Template Search

Template search with BLAST and HHBlits has been performed against the SWISS-MODEL template library (SMTL, last update: 2020-01-23, last included PDB release: 2020-01-17).

The target sequence was searched with BLAST against the primary amino acid sequence contained in the SMTL. A total of 134 templates were found.

An initial HHblits profile has been built using the procedure outlined in (Remmert et al.), followed by 1 iteration of HHblits against NR20. The obtained profile has then be searched against all profiles of the SMTL. A total of 635 templates were found.

##### Model Building

Models are built based on the target-template alignment using ProMod3. Coordinates which are conserved between the target and the template are copied from the template to the model. Insertions and deletions are remodelled using a fragment library. Side chains are then rebuilt. Finally, the geometry of the resulting model is regularized by using a force field. In case loop modelling with ProMod3 fails, an alternative model is built with PROMOD-II (Guex et al.).

##### Model Quality Estimation

The global and per-residue model quality has been assessed using the QMEAN scoring function (Benkert et al.) . For improved performance, weights of the individual QMEAN terms have been trained specifically for SWISS-MODEL.

##### Ligand Modelling

Ligands present in the template structure are transferred by homology to the model when the following criteria are met: (a) The ligands are annotated as biologically relevant in the template library, (b) the ligand is in contact with the model, (c) the ligand is not clashing with the protein, (d) the residues in contact with the ligand are conserved between the target and the template. If any of these four criteria is not satisfied, a certain ligand will not be included in the model. The model summary includes information on why and which ligand has not been included.

##### Oligomeric State Conservation

The quaternary structure annotation of the template is used to model the target sequence in its oligomeric form. The method (Bertoni et al.) is based on a supervised machine learning algorithm, Support Vector Machines (SVM), which combines interface conservation, structural clustering, and other template features to provide a quaternary structure quality estimate (QSQE). The QSQE score is a number between 0 and 1, reflecting the expected accuracy of the interchain contacts for a model built based a given alignment and template. Higher numbers indicate higher reliability. This complements the GMQE score which estimates the accuracy of the tertiary structure of the resulting model.

#### Table T1:

Primary amino acid sequence for which templates were searched and models were built.

```
MFVFLVLLPLVSSQCVNLTTRTQLPPAYTNSFTRGVYYPDKVFRSSVLHSTQDLFLPFFSNVTWFHAIHVSGTNGTKRFDNPVLPFNDGVYFASTEKSNI
IRGWIFGTTLDSKTQSLLIVNNATNVVIKVCEFQFCNDPFLGVYYHKNNKSWMESEFRVYSSANNCTFEYVSQPFLMDLEGKQGNFKNLREFVFKNIDGY
FKIYSKHTPINLVRDLPQGFSALEPLVDLPIGINITRFQTLLALHRSYLTPGDSSSGWTAGAAAYYVGYLQPRTFLLKYNENGTITDAVDCALDPLSETK
CTLKSFTVEKGIYQTSNFRVQPTESIVRFPNITNLCPFGEVFNATRFASVYAWNRKRISNCVADYSVLYNSASFSTFKCYGVSPTKLNDLCFTNVYADSF
VIRGDEVRQIAPGQTGKIADYNYKLPDDFTGCVIAWNSNNLDSKVGGNYNYLYRLFRKSNLKPFERDISTEIYQAGSTPCNGVEGFNCYFPLQSYGFQPT
NGVGYQPYRVVVLSFELLHAPATVCGPKKSTNLVKNKCVNFNFNGLTGTGVLTESNKKFLPFQQFGRDIADTTDAVRDPQTLEILDITPCSFGGVSVITP
GTNTSNQVAVLYQDVNCTEVPVAIHADQLTPTWRVYSTGSNVFQTRAGCLIGAEHVNNSYECDIPIGAGICASYQTQTNSPRRARSVASQSIIAYTMSLG
AENSVAYSNNSIAIPTNFTISVTTEILPVSMTKTSVDCTMYICGDSTECSNLLLQYGSFCTQLNRALTGIAVEQDKNTQEVFAQVKQIYKTPPIKDFGGF
NFSQILPDPSKPSKRSFIEDLLFNKVTLADAGFIKQYGDCLGDIAARDLICAQKFNGLTVLPPLLTDEMIAQYTSALLAGTITSGWTFGAGAALQIPFAM
QMAYRFNGIGVTQNVLYENQKLIANQFNSAIGKIQDSLSSTASALGKLQDVVNQNAQALNTLVKQLSSNFGAISSVLNDILSRLDKVEAEVQIDRLITGR
LQSLQTYVTQQLIRAAEIRASANLAATKMSECVLGQSKRVDFCGKGYHLMSFPQSAPHGVVFLHVTYVPAQEKNFTTAPAICHDGKAHFPREGVFVSNGT
HWFVTQRNFYEPQIITTDNTFVSGNCDVVIGIVNNTVYDPLQPELDSFKEELDKYFKNHTSPDVDLGDISGINASVVNIQKEIDRLNEVAKNLNESLIDL
QELGKYEQYIKWPWYIWLGFIAGLIAIVMVTIMLCCMTSCCSCLKGCCSCGSCCKFDEDDSEPVLKGVKLHYT
```

#### Table T2:

| Template | Seq Identity | Oligo-state | QSQE | Found by | Method | Resolution | Seq Similarity | Coverage | Description |
|---|---|---|---|---|---|---|---|---|---|
| 6acd.1.C | 76.47 | homo-trimer | 0.75 | HHblits | EM | NA | 0.54 | 0.93 | Spike glycoprotein |
| 6acc.1.A | 76.47 | homo-trimer | 0.74 | HHblits | EM | NA | 0.54 | 0.93 | Spike glycoprotein |
| 6acg.1.A | 76.47 | homo-trimer | 0.73 | HHblits | EM | NA | 0.54 | 0.93 | Spike glycoprotein |
| 6acg.1.B | 76.47 | homo-trimer | 0.73 | HHblits | EM | NA | 0.54 | 0.93 | Spike glycoprotein |
| 6acj.1.A | 76.47 | homo-trimer | 0.75 | HHblits | EM | NA | 0.54 | 0.93 | Spike glycoprotein |
| 6acj.1.B | 76.47 | homo-trimer | 0.75 | HHblits | EM | NA | 0.54 | 0.93 | Spike glycoprotein |
| 6acg.1.C | 76.47 | homo-trimer | 0.72 | HHblits | EM | NA | 0.54 | 0.93 | Spike glycoprotein |
| 6acj.1.C | 76.47 | homo-trimer | 0.74 | HHblits | EM | NA | 0.54 | 0.93 | Spike glycoprotein |
| 3jcl.1.A | 31.50 | homo-trimer | 0.38 | HHblits | EM | NA | 0.36 | 0.86 | Spike glycoprotein |
| 6b3o.1.A | 41.04 | homo-trimer | 0.24 | HHblits | EM | NA | 0.40 | 0.41 | Spike glycoprotein |
| 6b3o.1.A | 44.07 | homo-trimer | 0.23 | BLAST | EM | NA | 0.42 | 0.38 | Spike glycoprotein |
| 5x4s.1.A | 57.03 | monomer | - | BLAST | X-ray | 2.20Å | 0.49 | 0.21 | Spike glycoprotein |
| 3jcl.1.A | 31.82 | homo-trimer | 0.04 | BLAST | EM | NA | 0.37 | 0.14 | Spike glycoprotein |
| 3r4d.1.B | 22.32 | monomer | - | HHblits | X-ray | 3.10Å | 0.31 | 0.18 | Spike glycoprotein |
| 3r4d.2.B | 22.32 | monomer | - | HHblits | X-ray | 3.10Å | 0.31 | 0.18 | Spike glycoprotein |
| 1zva.1.A | 76.36 | homo-dimer | 0.02 | HHblits | X-ray | 1.50Å | 0.49 | 0.04 | E2 glycoprotein |
| 1wdf.1.A | 62.07 | homo-hexamer | - | BLAST | X-ray | 2.50Å | 0.47 | 0.05 | E2 glycoprotein |
| 1wdf.1.B | 62.07 | homo-hexamer | - | BLAST | X-ray | 2.50Å | 0.47 | 0.05 | E2 glycoprotein |
| 1wyy.1.A | 69.44 | monomer | - | BLAST | X-ray | 2.20Å | 0.48 | 0.06 | E2 Glycoprotein |
| 1wdf.1.A | 61.02 | homo-hexamer | - | HHblits | X-ray | 2.50Å | 0.47 | 0.05 | E2 glycoprotein |
| 1wdf.1.B | 61.02 | homo-hexamer | - | HHblits | X-ray | 2.50Å | 0.47 | 0.05 | E2 glycoprotein |
| 2beq.1.A | 76.32 | homo-trimer | 0.25 | HHblits | X-ray | 1.60Å | 0.49 | 0.03 | E2 GLYCOPROTEIN |

The table above shows the top 22 filtered templates. A further 477 templates were found which were considered to be less suitable for modelling than the filtered list.
5dfz.1.A, 1ztm.1.C, 3zrw.1.A, 6acg.1.C, 1g5g.1.A, 2j68.1.A, 4zs6.1.C, 3ghg.1.D, 3ghg.1.A, 6h7w.1.C, 3d0g.1.C, 6h7w.1.D, 4mms.1.B, 6h7w.1.I, 6h7w.1.J, 4w8n.1.B, 6gaj.1.B, 4jhw.1.C, 5vyh.1.A, 1bz4.1.A, 6acd.1.C, 2dd8.1.C, 5j3d.1.F, 4kqz.2.A, 1zvb.1.A, 6u7k.1.A, 1eq1.1.A, 5tok.1.C, 1br0.1.A, 5w9p.1.I, 3zbh.3.B, 3scj.1.B, 1wp8.1.A, 6gaj.1.C, 6nb3.1.G, 6nb3.1.A, 4i44.1.A, 3ja6.1.R, 3ja6.1.P, 6nzk.1.A, 3ja6.1.J, 3ja6.1.H, 6gap.1.C, 6gap.1.B, 6gap.1.A, 3ja6.1.L, 3lg7.1.A, 5yl9.1.B, 5yl9.1.A, 6t3f.1.A, 1wdf.1.A, 4mod.1.A, 6nb7.1.A, 6nb7.1.B, 6nb7.1.C, 2nrj.1.A, 1wyy.1.A, 3tul.3.A, 3dyt.1.A, 5x5b.1.C, 5wrg.1.A, 5x5b.1.A, 6s1k.1.G, 3tul.1.A, 6crx.1.B, 6crx.1.C, 6crx.1.A, 5k6b.1.A, 6u7h.1.A, 6tys.1.A, 5xgr.7.A, 5nen.1.A, 5ejb.1.A,

5nen.1.B, 3zbh.1.A, 3zbh.1.B, 5zvk.1.E, 5zvk.1.D, 5zvk.1.F, 5zvk.1.A, 5zvk.1.C, 3zbh.3.A, 4wsg.1.A, 4wsg.1.B, 4wsg.1.C, 3rrr.1.F, 3rrr.1.D, 5c69.1.A, 3rrr.1.B, 6fki.1.D, 5k6h.1.A, 6c6z.1.A, 6nyx.5.B, 5yzc.1.B, 6qaj.1.B, 6qaj.1.A, 3euh.1.B, 3euh.1.A, 5e5w.1.B, 3maw.1.A, 5x4s.1.A, 5e30.1.D, 6crw.1.A, 6crw.1.C, 6crw.1.B, 6qfy.1.A, 1z56.1.B, 5xlr.1.A, 6eag.1.A, 5xjk.1.A, 6nyx.1.A, 4h14.1.A, 1zv7.1.A, 1zv7.1.B, 6s7o.1.E, 6ead.1.A, 2vs0.1.A, 3u0c.1.A, 6gaj.1.A, 3gvm.2.A, 5w9a.1.D, 4yy9.1.B, 5w9a.1.A, 4ilo.1.A, 3rpu.1.A, 3dyu.2.A, 5w9i.1.J, 2lfs.1.A, 5xgr.1.A, 1t98.1.A, 1t98.1.B, 4zpt.2.B, 5e64.1.B, 4jeu.1.B, 4zpw.1.A, 6fln.1.B, 6fln.1.A, 5m4y.2.A, 5zvm.1.B, 5zvm.1.A, 5udd.1.C, 5udd.1.B, 5udd.1.A, 5zvm.1.D, 3sci.1.B, 5k6g.1.A, 5cws.2.E, 5do2.2.C, 3scl.1.B, 6cv0.1.A, 4l3n.1.A, 5w9h.1.L, 3d0h.1.C, 6ous.2.F, 6ous.2.D, 6ous.2.B, 4uxv.1.A, 6j11.1.A, 4kr0.1.D, 6eam.1.A, 5zxv.1.A, 2ghv.1.A, 5ude.1.A, 6dc5.3.A, 3g67.1.B, 2bez.1.A, 2bez.1.B, 3g67.1.A, 2fyz.1.E, 2fyz.1.A, 5yy5.1.A, 5jhf.2.C, 6dc5.1.A, 2xz3.1.A, 5w23.1.A, 5w23.1.B, 5cws.1.E, 1wnc.1.A, 1wnc.1.B, 1wnc.1.C, 1wnc.1.D, 1wnc.1.E, 1wnc.1.F, 6n9h.1.A, 4cg4.3.A, 4cg4.3.B, 6qu1.1.A, 6ack.1.B, 6ack.1.C, 4cfg.1.B, 4cfg.1.A, 4cg4.1.B, 4cg4.1.A, 1zv8.2.D, 2ch7.1.B, 6grk.1.A, 4cg4.2.A, 5x58.1.A, 4i3m.1.A, 5w9m.1.A, 3n27.1.B, 3n27.1.A, 5w9m.1.D, 6jx7.1.A, 2rai.1.A, 5yxw.1.B, 5x5f.1.C, 6cxc.1.H, 4hpq.1.C, 4mmu.1.B, 4zyp.1.A, 4zyp.1.C, 2rak.1.A, 5ijn.1.R, 6nb6.1.A, 5ijn.1.H, 5ijn.1.L, 5ijn.1.G, 5ijn.1.F, 6eaf.1.A, 5lg4.1.A, 2j69.1.A, 3jcl.1.A, 5tpn.1.A, 1wp7.1.C, 1wp7.1.A, 4ut1.1.A, 4fzs.1.B, 4fzs.1.A, 3zrv.1.B, 1m1j.1.D, 3zrv.1.A, 1m1j.1.A, 1svf.1.A, 4kqz.1.A, 2b9b.1.A, 2b9b.1.B, 2b9b.1.C, 5kuc.1.A, 5e2z.1.B, 4nqj.1.B, 4nqj.1.C, 4nqj.1.A, 6ntx.1.A, 1z56.1.A, 5l1x.1.D, 5l1x.1.F, 5l1x.1.B, 3zrw.2.A, 3gwk.1.A, 3gwk.1.B, 3g6b.1.B, 3g6b.1.A, 1htm.1.F, 1htm.1.D, 1htm.1.B, 1fio.1.A, 6q04.1.A, 4l72.2.A, 2ghw.1.A, 6naf.1.A, 6nro.1.A, 3tul.4.A, 3tul.2.A, 5e32.1.B, 6ack.1.A, 3bgf.1.A, 5k6c.1.A, 6nb6.1.C, 6nb6.1.B, 3kpe.1.A, 5toj.1.A, 2ieq.1.A, 5toj.1.C, 5toj.1.B, 5zhy.1.A, 5zhy.1.C, 6bfu.1.A, 3ghg.2.A, 3dyu.1.A, 3ghg.2.D, 4cjd.1.A, 4cgk.1.A, 4cgk.1.B, 2ajf.1.B, 6b7n.1.A, 4njl.1.A, 3ci9.1.A, 3ci9.1.B, 4mmt.1.B, 3zcc.1.A, 3zcc.1.B, 5m4y.1.A, 5x5c.1.B, 5x5c.1.C, 5x5c.1.A, 2ba2.1.C, 2beq.1.F, 2beq.1.E, 2beq.1.D, 2beq.1.A, 4dag.1.A, 1ruy.1.B, 4akv.1.B, 2dnx.1.A, 4akv.1.A, 5tdg.1.A, 3lnr.1.A, 5tdg.1.C, 5tdg.1.B, 1ztm.1.A, 6acg.1.A, 6acg.1.B, 1ztm.1.B, 4xak.1.A, 5c6b.1.A, 6h2f.1.F, 6h2f.1.B, 6h2f.1.A, 5m4y.3.A, 4tn3.1.A, 4tn3.1.B, 5i08.1.A, 3zbh.4.A, 3zbh.4.B, 2lfs.1.B, 1g2c.1.A, 5e30.1.B, 5w9k.1.L, 6gao.1.B, 6gao.1.C, 5w9k.1.K, 1wdg.1.B, 1wdg.1.A, 5k6i.1.A, 5k6f.1.A, 6oe5.1.A, 5w9j.1.D, 5w9j.1.L, 6grk.3.A, 4mmv.1.B, 6fkf.1.I, 5w9m.1.E, 3ja6.1.N, 3zbh.2.B, 3zbh.2.A, 6mjz.1.B, 6mjz.1.C, 6mjz.1.A, 6f1t.1.h, 1zv8.1.B, 3c98.1.B, 1zv8.1.A, 1zv8.1.D, 1zv8.1.E, 2q12.1.A, 6apd.1.C, 3rrt.1.B, 6apd.1.A, 3rrt.1.D, 6j9r.1.A, 6j9r.1.B, 6eaj.1.A, 3u0c.2.A, 4zpw.2.A, 6cs2.1.C, 6cs2.1.B, 6cs2.1.A, 3rrt.1.F, 6f1t.1.U, 6ous.1.B, 6ous.1.D, 6ous.1.F, 5zvk.1.B, 6h3a.1.B, 6h3a.1.D, 2w6d.1.B, 6pxh.1.A, 2w6d.1.A, 5zuv.1.A, 5zuv.1.C, 5zuv.1.B, 6crv.1.A, 6eal.1.A, 1wdf.1.B, 6q0s.1.A, 5ea3.1.A, 5x4r.1.A, 6c6y.1.C, 5gmq.1.A, 6dc5.2.A, 5szs.1.A, 4p1w.1.F, 4p1w.1.C, 5tdl.1.A, 5xbj.1.A, 4l3n.2.A, 4lws.2.A, 6l8q.1.B, 3zx6.1.A, 3zx6.1.B, 6s1k.1.P, 1mg1.1.A, 6s1k.1.H, 6s1k.1.I, 6s1k.1.J, 6s1k.1.L, 6s1k.1.M, 6s1k.1.N, 6s1k.1.O, 4gip.1.D, 4gip.1.F, 6s1k.1.E, 6s1k.1.F, 4gip.1.B, 5evm.1.A, 4cg4.2.B, 5evm.1.C, 4zpt.1.C, 5e2z.1.F, 1g2c.4.C, 5e2z.1.D, 5jhf.1.F, 3d0i.1.C, 3sck.1.B, 2raj.1.A, 4jeh.1.B, 2fxp.1.A, 3rpu.1.D, 5ea4.1.A, 2vrz.1.A, 2vrz.1.B, 4qzv.1.B, 6acj.1.A, 6acj.1.C, 6acj.1.B, 3tnu.1.B, 3tnu.1.A, 6dc3.1.C, 4ilo.2.A, 5zvm.1.F, 6pz8.1.A, 6gao.1.A, 6pz8.1.B, 5gyq.1.A, 6eah.1.A, 5huf.1.B, 4mmr.1.B, 5w9n.1.G, 6ean.1.A, 6eai.1.A, 6acc.1.A, 1zva.1.A, 6eae.1.A, 5y05.1.A, 4abx.1.B, 2ba2.1.A, 2ba2.1.B, 4abx.1.A, 5kwb.1.A, 5udc.1.I, 5wb0.1.A, 4rsi.1.B, 4yy0.1.F, 4yy0.1.D, 4yy0.1.B, 5u68.1.A, 5u68.1.C, 5u68.1.B, 6jhy.1.A, 6grj.1.A, 6grj.1.B, 5w9l.1.H, 5e35.1.B, 3oja.1.B, 5w9l.1.B, 5w9l.1.E
